# VCFShark: how to squeeze a VCF file

**DOI:** 10.1101/2020.12.18.423437

**Authors:** Sebastian Deorowicz, Agnieszka Danek

## Abstract

**Summary:** The VCF files with results of sequencing projects take a lot of space. We propose VCFShark squeezing them up to an order of magnitude better than the de facto standards (gzipped VCF and BCF).

**Availability and Implementation:** https://github.com/refresh-bio/vcfshark

**Supplementary information:** Supplementary data are available at publisher’s Web site.

## 1 Introduction

Accessibility of cheap sequencing technologies allowed comparative genomics to extend its field of interests from viruses in the 1980s (Argos *et al*., 1984) to such complex species like humans in the 2010s (Sudmant *et al*., 2015). Nowadays, the largest sequencing projects cover tens or even hundreds of thousands of individuals (McCarthy *et al*., 2016; Bycroft *et al*., 2018). It seems obvious that in the near future we will see collections of millions of human genomes.

Genome collections are usually stored in the Variant Call Format (VCF) (Danecek *et al*., 2011) files. It is composed of a series of lines, each representing a description of a single variation. The first part of each line contains mandatory fields, like a chromosome, coordinates, reference allele, alternative alleles, etc. It is followed by an unlimited number of optional fields, that can be, e.g., genotypes, aggregate data, like alternate allele frequency, statistics for various (sub)populations.

VCF files can be huge, so their storage, maintenance, transfer is challenging and data compression is a must. Nowadays, they are gzip-compressed or stored in BCF files (internally gzip-compressed). Nevertheless, often much more can be done. Some partial solutions (focusing just on genotypes) are PBWT (Durbin, 2014), BGT (Li, 2015), GTC (Danek and Deorowicz, 2018), GTShark (Deorowicz and Danek, 2019). Recently, Lan *et al*. (2020) proposed genozip, a VCF-specialized compressor.

In this article, we present VCFShark—a dedicated fully-fledged compressor of VCF files. It significantly outperforms the universal tools in terms of compression ratio; sometimes its advantage is severalfold. Usually, it is also significantly better than genozip. At the same time, the compression speeds are similar to the competitors.

## 2 Methods

VCFShark processes the input file in a column-wise manner. Below, we give a rough description of the algorithm and more details can be found in Supplementary Section 1. The fields CHROM, POS, ID, REF, ALT, QUAL, FILTER are processed as separate *fixed* streams. The INFO field can contain a number of subfields and they are distributed into separate *info* streams. The FORMAT field, together with related sample-specific data, is also split into separate *format* streams. A special treatment is implemented for the *genotype* stream.

All streams are split into *chunks* of size 8 MB (256 MB for *genotype*). The processing of chunks is partially independent. The *fixed* chunks are compressed by a BSC compressor (http://libbsc.com/) as they are, except for the *pos* stream, which is firstly delta-coded. The “narrow” textual *info* and *format* streams (i.e., the average length of the subfield is less than 64 bytes) are also independently BSC-compressed.

The “wider” textual chunks (often containing the majority of VCF data) are firstly preprocessed. We identify common substrings, construct an adaptive dictionary (i.e., growing during the processing of the file) of them, and replace them in the text by their unique identifiers. Moreover, we identify numbers and store them as binary values. Finally, we use the run-length encoding of a series of 0s and ‘|’s, as well as use some minor tricks. Such preprocessed chunks are then BSC-compressed. The dictionary is for a stream, so the processing of such chunks is not fully independent.

The streams with numeric data (integer and real values) are treated in a special way. We identify the “pattern” of no. of values, e.g., each subfield contains 0 or 1 value, each line contains the same number of values. Then we estimate the entropy (order-0, order-1, or order-2) taking as a context various previously encoded numbers, e.g., the previous in this line, the one at the same position in the previous line. We select the order and the context construction method which minimizes the entropy estimation. Then, we process the numeric values using an entropy coder (i.e., range coder).

For handling genotypes we extended GTShark algorithm (as well as the generalized positional Burrows–Wheeler transform it is based on) to support VCF files with non-uniform ploidy, present for example in human sex chromosomes.

Various streams contain related values, e.g., the same value in separate streams, the same number of integers in separate streams. Therefore, we implemented algorithms discovering the functional dependencies between streams. Thanks to this, some streams can be described in a very compact way as a function of the other stream.

## 3 Results

For evaluation, we used 11 datasets (characterized in Supplementary Section 3) from large sequencing projects. They differ in size and contents. For example, HRC contains 27,165 human genotypes at 40.4M variants. Some others contain just aggregate data, i.e., without genotypes. Rice is a collection of gVCF files, each containing just a single individual data.

For comparison we used BCF (binary version of VCF), pigz (parallel gzip, commonly used in the field), 7z (one of the best universal compressors), and genozip (modern VCF files compressor). The summary of the results is given in Figure 1, while the details can be found in Supplementary Worksheet 1. The compressors were configured to use 8 threads.

**Fig. 1.**
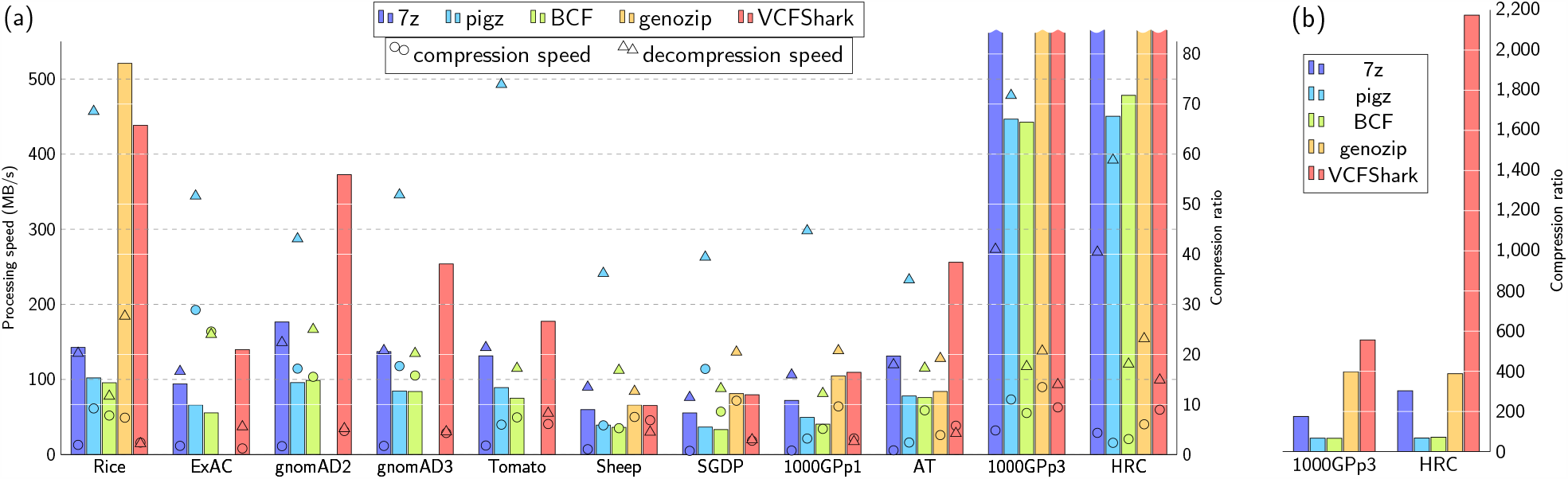
Results of compression of 11 sets of VCF files using 5 different tools. (a) Compression ratio (vertical bars), compression and decompression speed (circe and triangle markers) for each tool and each dataset. To highlight the differences between the ratios on all sets, the compression ratios larger than 85 (achieved for 1000GPp3 and HRC sets) are cut off (see plot (b) for complete ratio results). Genozip does not support the compression of some datasets (Tomato, ExAC, gnomAD2, gnomAD3). (b) Compression ratios for 1000GPp3 and HRC datasets.

It is easy to notice that VCFShark offers the best compression ratios for most of the datasets. Its advantage over the competitors depends strongly on the contents of the dataset. When the majority of VCF is genotype data (HRC, 1000GPp3), VCFShark dominates over BCF, pigz, and 7z by a large margin, achieving 3-to 32-fold better compression. It is mainly a result of an algorithm for compression of genotypes. The advantage over genozip, which uses similar compression for genotypes (also based on GTShark), is also significant, up to 5.5-fold for HRC.

For the datasets with genotypes supported by other (mainly numeric) data (AT, 1000GPp1, Sheep, Tomato, SGDP) the gain over 7z is from 9% to 95% and even larger over BCF and pigz. The gain over genozip is 3-fold for AT and 5% for 1000GPp1, while for sets with fewer genotypes per line, the results are similar, up to 2% worse for VCFShark.

For aggregate datasets (ExAC, gnomAD2, gnomAD3) the gain over second best, 7z, is from 48% to 111 %. The gVCF files (Rice) are much easier to compress by VCFShark than by 7z (almost 3-fold advantage), but Genozip achieves a 16% better compression ratio than VCFShark.

In compression, VCFShark is usually a few times faster than 7z, but in decompression, it is a few times slower. Genozip is usually faster than VCFShark, but does not support all VCF files. Nevertheless, the absolute operating speeds of VCFShark between 15 MB/s and 100 MB/s (in most cases) should be acceptable in typical scenarios.

The main memory used by the compressors varies, in most cases, from single MBs (gzip and BCF), by 1–3 GB (Genozip), to 1–5 GB for VCFShark. There are, however, datasets like Sheep and aggregate ones that require more memory, e.g., 8 GB by genozip and 12–50 GB by VCFShark.

## 4 Conclusions

We proposed a novel compression algorithm for VCF files. Our main goal was to develop a practical tool that will be able to squeeze the data a few times better than pigz. For the largest dataset, with a lot of genotype data, its advantage is more than 30-fold. The advantage over genozip (the only VCF-specialized compressor) is usually significant, however, sometimes the competitor is a bit better. The good compression ratios and (de)compression speeds should make VCFShark suitable for applications where huge VCF files must be stored or transmitted.

## Supporting information

Supplementary Section 1

Supplementary Section 3

## Funding

The work was supported by National Science Centre, Poland, project DEC-2017/25/B/ST6/01525 and by POIG.02.03.01-24-099/13 grant: “GeCONiI—Upper Silesian Center for Computational Science and Engineering” (the infrastructure).

## Conflict of Interest

none declared.

## Notes

https://github.com/refresh-bio/VCFShark

### Competing Interest Statement

The authors have declared no competing interest.

