## Supplementary Section 1 for "VCFShark: how to squeeze a VCF file"

### Supplementary material for article:
### VCFShark:

Sebastian Deorowicz Agnieszka Danek

December 14, 2020

**4 Environment 16**

**5 Additional results 17**

### 1 Methods

#### 1.1 VCF file characteristics

VCF files are textual and are composed of two parts: *header* and *data* [2]. The *header* part describes the contents of the file, e.g., specifies the subfields in some fields of the *data* part. The *data* part is composed of a series of lines describing successive variants. Each line is composed of the following fields/columns:

1. CHROM — identifier of the chromosome,
2. POS — position of the variant at the chromosome,
3. ID — identifier of the variant,
4. REF — reference base(s),
5. ALT — alternate base(s); a list of alternatives can be given here,
6. QUAL — quality,
7. FILTER — filter status,
8. INFO — additional information; a list of pairs: subfield name – subfield value; the valid subfields are defined in the header; the list can be empty or exceedingly long,
9. FORMAT specification (optional) — list and order of value types for the sample-specific data,
10. FORMAT data (optional, one for each sample) — genotypes (GT) and other sample-specific data; the number of subfields (for each sample) is defined in the FORMAT specification of the variant; the number of fields of this type is equal to the number of samples.

The first 8 fields are mandatory. They describe a single variant. The first 7 fields are fixed in the sense that they usually contain a single or a small number of values. There can be some exceptions (e.g., multiple alternate alleles or a list of identifiers), but they are rare. When the VCF file contains descriptions of many samples, the size of the fixed fields is usually a small fraction of the size of the whole VCF file.

#### 1.2 Splitting of VCF files into streams and chunks

VCFShark splits the VCF file into many separate *streams*. Each fixed field is treated as a separate *fixed* stream. INFO field is split into as many *info* streams as the number of INFO subfields defined in the file header. Similarly, FORMAT column is split into as many *format* streams as the number of FORMAT subfields defined in the header. Therefore, all sample-specific data for a subfield go to the same stream. The genotype (GT) data are treated in a special way and we denote its stream as *genotype*.

The streams are then divided into *chunks*. The size of most chunks is almost 8 MB (we complete a chunk if adding new values would break the limit). The only difference is for *genotype* chunks that are of size up to 256 MB.

#### 1.3 Compression of sizes of fields for each variant

The data from successive lines are concatenated in chunks, so we store an array of 32-bit integers describing the size of the field or subfield in each variant. Thus, for each stream we store two compressed sub-streams. First, for the sizes of the (sub)field, i.e., *stream-sizes*. Second, for the data in the (sub)field, i.e., *stream-data*.

The *stream-sizes* sub-streams are compressed using the BSC compressor (`http://libbsc.com/`), a universal tool based on the Burrows–Wheeler transform. If not stated explicitly, in the rest of the section we focus on the compression of the *stream-data* sub-streams.

##### 1.4 Compression of header and *fixed* streams

The header as well as the chunks from streams CHROM, ID, REF, ALT, QUAL, FILTER are compressed separately with a use of the BSC compressor. Since the variants are usually ordered according to CHROM:POS, the differences between the values in the POS stream are small. Therefore, these values are delta-coded, i.e., the differences between successive numbers are stored, which helps the compression. The delta-coded values are then compressed using the BSC tool.

##### 1.5 Compression of "narrow" textual *info* and *format* streams

We classify the textual *info* and *format* streams as "narrow" if the average length of the field in each chunk line is less than 64 bytes. The remaining textual streams are classified as "wide". The "narrow" streams are compressed using the BSC algorithm.

##### 1.6 Compression of "wide" textual *info* and *format* streams

The "wide" streams (if present) usually contribute to the total size of the VCF files significantly. Therefore, we decided to prepare a specialized compression algorithm for them.

For a stream we maintain two dictionaries: *candidate* and *final*. The processing of each chunk is as follows. First, we split the chunk into a list of tokens. The token types are:

- *word* — sequence of symbols from the alphabet $\{\texttt{A}, \ldots, \texttt{Z}, \texttt{a}, \ldots, \texttt{z}, \_, \texttt{0}, \ldots, \texttt{9}, (, ), \&, /\}$ (starting from a symbol from the alphabet $\{\texttt{A}, \ldots, \texttt{Z}, \texttt{a}, \ldots, \texttt{z}, \_\}$) of minimal length 6,
- *bar-run* — sequence of '|' characters,
- *0-run* — sequence of '0' characters,
- *number* — sequence of decimal digits ('0', ..., '9') not starting from '0',
- *base* — one of sequences 'A:', 'C:', 'G:', 'T:',
- *other* — single symbol not matching the criteria to be classified as other token type.

For each *word* token we check whether it is present in the *final* dictionary. If not, we add it to the *candidate* dictionary or increment the counter related to it in the *candidate* dictionary. If the related counter becomes 16, we move the word from the *candidate* to the *final* dictionary and assign a code to it (size of the *final* dictionary before adding the new *word*).

Now we are ready to transform the chunk. First of all, we output the words inserted to the *final* dictionary while processing of the current chunk. Then, we process the chunk token by token and for:

- *word* — store the word in plain if it is not in the *final* dictionary; otherwise store:
  - byte `0xff` followed by 1-byte code (for codes $< 256$),
  - byte `0xfe` followed by 2-byte code (for codes $< 256^2 + 256$),
  - byte `0xfd` followed by 3-byte code (for codes $< 256^3 + 256^2 + 256$),

  we do not allow the dictionary to be larger than $256^3 + 256^2 + 256 - 1$,
- *bar-run* — store the length using byte values from `0xed` to `0xfc`,
- *0-run* — store the length using byte values from `0xe3` to `0xed`,
- *number* — store the number in binary using byte values from `0x80` to `0xe3`; numbers longer than 15 digits are encoded in plain,
- *base* — store byte `0x01` for 'A:', `0x02` for 'C:', `0x03` for 'G:', `0x04` for 'T:',
- *other* — store byte in plain.

The output of the preprocessing is compressed using the BSC tool.

##### 1.7 Compression of numeric *info* streams

For each chunk, the first stage is to determine the pattern of sizes of subfields. The result of this can be:

- *one* — the size is 1 for each variant (i.e., there is a single number at the subfield for each variant),
- *zero-one* — the size is 0 or 1 for each variant,
- *const* — the size is larger than 1 but the same for each variant,
- *zero-const* — the size is 0 or some constant number (larger than 1) for each variant,
- *other* — something not matching the above-listed criteria.

If the pattern is the same as for the previous chunk, we assume that the context construction will also be the same. Otherwise (and of course also for the first chunk) we determine the best context construction method. To this end, we estimate the entropy for various methods of construction of a context. We try contexts composed of 0, 1, or 2 symbols. The symbols we try depend on the pattern of sizes:

- *one* or *zero-one* — for the $i$-th variant we pick the codes for numbers (how we determine the code for the numbers will be described in the next paragraph) from the $(i-1)$-th and $(i-2)$-th variant (in case of *zero-one* we pick one or two recently seen numbers),
- *const* or *zero-const* — for the $i$-th variant we try nine possibilities; for simplicity let us assume the *const* pattern and the notation $v_i^j$ for the value in the $i$-th variant at the $j$-th position; the context can be:
  - empty,
  - $v_i^{j-1}$,
  - $v_i^{j-2}$, $v_i^{j-1}$,
  - $v_{i-1}^j$,
  - $v_{i-2}^j$, $v_{i-1}^j$,
  - $v_i^{j-1}$, $v_{i-1}^j$,
  - $j$, $v_i^{j-1}$,
  - $j$, $v_{i-1}^j$,
  - $j$.

  The strategy for *zero-const* is similar, we simply ignore the "empty" variants,
- *other* — we treat the sequence of numbers exactly as for *one* pattern ignoring that the situation is different; handling this pattern in a special way was left for the future.

Instead of encoding the numbers as they are, we use an adaptive dictionary (of max. size $256^3$). When we see the value for the first time we encode it in plain and add it to the dictionary (with the code equal to the dictionary size before inserting the number). Otherwise we store the code. We use the context determined as described above, and expand it by the already encoded bytes of the code.

#### 1.8 Compression of numeric *format* streams

Similarly like for the *info* streams, we determine the pattern of sizes of subfields. Here the results can be:

- *one* — the number of values for each variant is equal to the number of samples,
- *many* — otherwise (the number of values for each variant must be a multiplicity of the number of samples).

Now the processing depends on the pattern. Here we also encode the never-seen-before numbers in plain, but for the rest we use codes (similar as described for the *info* numeric values) and proceed as follows. For the pattern:

- *one* — we use the context formed of: the value in the left-neighbor sample in the same variant and the value in the previous variant for the current sample,
- *many* — we use the context formed of: the position of the value in the subfield, two recently processed codes for this variant; if this is the first value of the subfield we use also the sample number.

In the *many* mode, we also check whether the complete subfield for the sample (composed of a few numbers) is the same as in the previous variant for the same sample. If so, we encode just such an information.

##### 1.9 Compression of *genotype* stream

The *genotype* stream is encoded with a modified version of our former GTShark compressor of genotype data [3]. This algorithm was extended to support change of ploidy in the VCF file. Such a situation can happen for example for human sex chromosomes. The modification is quite simple. When we notice that the ploidy of the current variant is different than for the previous variant, we just expand (or reduce) the column indices in the internal gPBWT (generalized positional Burrows–Wheeler transform) algorithm, which is a core of the GTShark algorithm. The gPBWT is a generaization of PBWT [5], which was introduced in [4]. This is a different approach than used by genozip [8], which expands the genotypes in the whole chunk of VCF files to the largest ploidy present in this chunk.

##### 1.10 Exploring of functional dependencies between columns

In the final stage, after the main compression, VCFShark tries to reduce the compressed file even more. We implemented two strategies.

The first (simpler) is to compare the compressed data for the separate sub-streams (*stream-size*, *stream-data*). If we notice something identical, we remove one of the sub-streams. Such a strategy is quite fast and for some files can reduce the size of the compressed file by a few percent.

The second is to trace the functional dependencies between values of *info* or *format* data in separate streams. Such dependencies should be observed for example for *info* streams AC (allele count) and AF (allele frequency). Therefore, after processing of the whole file we are able to compare the space necessary to describe the "function" to the size of the compressed streams and store the "function" if it is beneficial. This worked, i.e., we observed some (not so big, however) gains. Unfortunately, the processing times, especially for files with large number of streams, were a few times longer. Therefore, we decided to disable this feature in the current version of the software and leave the investigations on this approach for the future work.

##### 1.11 Implementation

The implementation was prepared in the C++14 language and is distributed under the GNU GPL 3 license. It is parallelized to make use of modern multicore CPUs. The parallelization is at the level of streams, so the scalability depends on the contents of the input file.

#### 2 Examined programs

The following programs were used in the experimental part. The running parameters are also given.

- pigz v. 2.2.3:
  - Compression:
    pigz -9 -p 8 <in_file>
  - Decompression:
    pigz -9 -d <in_file>
- bcftools v. 1.4:
  - Compression:
    bcftools view -Ob -l9 --threads 8 -o <out_file> <in_file>
  - Decompression:
    bcftools view -Ov --threads 8 -o <out_file> <in_file>
- 7z v. 16.02:
  - Compression:
    7za a -mx9 -mmt8 <out_file> <in_file>
  - Decompression:
    7za e <in_file>
- genozip v. 8.0.3:
  - Compression:
    genozip -i vcf --gtshark -@ 8 -o <out_file> <in_file>
  - Decompression:
    genounzip -@ 8 -o /<out_file> <in_file>
- VCFShark v. 1.0:
  - Compression:
    vcfshark compress -t 8 <in_file> <out_file>
  - Decompression:
    vcfshark decompress -t 8 <in_file> <out_file>

### 3 Datasets

This section presents datasets used in the experimental part. Table 1 presents a summary characterisation of datasets.

Table 1: Characterisation of datasets.

| Dataset | Size (GB) | Files (number of) | Samples (number of) | Variant sites (number of) | FILTER | INFO | FORMAT | GT present |
|---|---|---|---|---|---|---|---|---|
| | | | | | (number of possible fields) | | | |
| HRC | 4,305 | 25[a] | 27,165 | 40,405,505 | 1 | 4 | 1 | + |
| 1000GPp3 | 854 | 25[a] | 2,504 | 84,805,772 | 1 | 27[d] | 1[d] | + |
| 1000GPp1 | 900 | 26[a] | 1,092 | 39,728,277 | 1 | 22[d] | 3[d] | + |
| AT | 1,287 | 5[a] | 1,135 | 119,146,348 | 3 | 1 | 3 | + |
| Tomato | 12 | 1 | 361 | 11,620,517 | 1 | 9 | 3 | + |
| Sheep | 85 | 1 | 99 | 42,438,841 | 1 | 26 | 7 | + |
| SGDP | 28 | 26[b] | 26 | 110,291,925 | 2 | 23 | 6 | + |
| ExAC | 41 | 1 | 60,706[c]. | 9,362,318 | 17 | 97 | 0[e] | - |
| gnomADv2 | 3,490 | 23[a] | 15,708[c] | 261,942,336 | 4 | 568 | 0 | - |
| gnomADv3 | 1,961 | 24[a] | 71,702[c] | 707,950,943 | 4 | 157 | 0 | - |
| Rice | 164 | 6[b] | 6 | 20,617,610,298 | 2 | 20 | 5 | + |

[a]single file describe single chromosome; [b]single file describe single individual; [c]aggregate VCF files with no genotype data about individuals; [d]in majority of files; [e]7 possible fields defined in the header, but zero used, as there is no FORMAT fields.

#### 3.1 1000GPp1

##### Description

The dataset describes a total of 39, 728, 277 variant sites at 1, 092 *H. sapiens* individuals.

It consists of 26 VCF files, corresponding to 22 autosomes, chromosomes X, Y (two files) and MT.

The number of allowed FILTER/INFO/FORMAT fields (based on the headers of the files):
- autosomes and chr X: 26 (1/22/3), including GT in FORMAT,
- chr Y ("phase1"): 24 (1/17/6), including GT in FORMAT,
- chr Y ("genome_strip"): 35 (1/28/6), including GT in FORMAT,
- chr MT: 4 (1/2/1), including GT in FORMAT.

##### Source

The dataset is from the Phase 1 of the 1000 Genomes Project [12].

The files were downloaded from:
`ftp://ftp.1000genomes.ebi.ac.uk/vol1/ftp/phase1/analysis_results/integrated_call_sets/`

The complete list of compressed VCF files used:

```
ALL.chr1.integrated_phase1_v3.20101123.snps_indels_svs.genotypes.vcf.gz
ALL.chr2.integrated_phase1_v3.20101123.snps_indels_svs.genotypes.vcf.gz
ALL.chr3.integrated_phase1_v3.20101123.snps_indels_svs.genotypes.vcf.gz
ALL.chr4.integrated_phase1_v3.20101123.snps_indels_svs.genotypes.vcf.gz
ALL.chr5.integrated_phase1_v3.20101123.snps_indels_svs.genotypes.vcf.gz
ALL.chr6.integrated_phase1_v3.20101123.snps_indels_svs.genotypes.vcf.gz
ALL.chr7.integrated_phase1_v3.20101123.snps_indels_svs.genotypes.vcf.gz
ALL.chr8.integrated_phase1_v3.20101123.snps_indels_svs.genotypes.vcf.gz
ALL.chr9.integrated_phase1_v3.20101123.snps_indels_svs.genotypes.vcf.gz
ALL.chr10.integrated_phase1_v3.20101123.snps_indels_svs.genotypes.vcf.gz
```

ALL.chr11.integrated_phase1_v3.20101123.snps_indels_svs.genotypes.vcf.gz
ALL.chr12.integrated_phase1_v3.20101123.snps_indels_svs.genotypes.vcf.gz
ALL.chr13.integrated_phase1_v3.20101123.snps_indels_svs.genotypes.vcf.gz
ALL.chr14.integrated_phase1_v3.20101123.snps_indels_svs.genotypes.vcf.gz
ALL.chr15.integrated_phase1_v3.20101123.snps_indels_svs.genotypes.vcf.gz
ALL.chr16.integrated_phase1_v3.20101123.snps_indels_svs.genotypes.vcf.gz
ALL.chr17.integrated_phase1_v3.20101123.snps_indels_svs.genotypes.vcf.gz
ALL.chr18.integrated_phase1_v3.20101123.snps_indels_svs.genotypes.vcf.gz
ALL.chr19.integrated_phase1_v3.20101123.snps_indels_svs.genotypes.vcf.gz
ALL.chr20.integrated_phase1_v3.20101123.snps_indels_svs.genotypes.vcf.gz
ALL.chr21.integrated_phase1_v3.20101123.snps_indels_svs.genotypes.vcf.gz
ALL.chr22.integrated_phase1_v3.20101123.snps_indels_svs.genotypes.vcf.gz
ALL.chrX.integrated_phase1_v3.20101123.snps_indels_svs.genotypes.vcf.gz
ALL.chrY.phase1_samtools_si.20101123.snps.low_coverage.genotypes.vcf.gz
ALL.chrY.genome_strip_hq.20101123.svs.low_coverage.genotypes.vcf.gz
ALL.chrMT.phase1_samtools_si.20101123.snps.low_coverage.genotypes.vcf.gz

###### Comments

The header of the ALL.chrY.genome_strip_hq.20101123.svs.low_coverage.genotypes.vcf file was modified, to comply with the VCF specification, i.e.:

- The “GL” field was declared as “Number=G”, insted of “Number=.”
- The missing definition of “SOURCE” field was added, assuming type String (INFO field): ##INFO=<ID=SOURCE,Number=1,Type=String>
- The missing definition of “FT” field was added, assuming type String (FORMAT field): ##FORMAT=<ID=FT,Number=1,Type=String>

##### 3.2 1000GPp3

###### Description

The dataset describes a total of 84,805,772 variant sites at 2,504 *H. sapiens* individuals.

It consists of 25 VCF files, corresponding to 22 autosomes, chromosomes X, Y and MT.

The number of allowed FILTER/INFO/FORMAT fields (based on the headers of the files):

- autosomes: 29 (1/27/1), including GT in FORMAT,
- chr X: 30 (1/28/1), including GT in FORMAT,
- chr Y: 32 (7/16/9), including GT in FORMAT,
- chr MT: 5 (2/2/1), including GT in FORMAT.

###### Source

The dataset is from the Phase 3 of the 1000 Genomes Project [14].

The files were downloaded from:
ftp://ftp.1000genomes.ebi.ac.uk/vol1/ftp/release/20130502/

The complete list of compressed VCF files used:

ALL.chr1.phase3_shapeit2_mvncall_integrated_v5a.20130502.genotypes.vcf.gz
ALL.chr2.phase3_shapeit2_mvncall_integrated_v5a.20130502.genotypes.vcf.gz
ALL.chr3.phase3_shapeit2_mvncall_integrated_v5a.20130502.genotypes.vcf.gz
ALL.chr4.phase3_shapeit2_mvncall_integrated_v5a.20130502.genotypes.vcf.gz
ALL.chr5.phase3_shapeit2_mvncall_integrated_v5a.20130502.genotypes.vcf.gz
ALL.chr6.phase3_shapeit2_mvncall_integrated_v5a.20130502.genotypes.vcf.gz
ALL.chr7.phase3_shapeit2_mvncall_integrated_v5a.20130502.genotypes.vcf.gz

```
ALL.chr8.phase3_shapeit2_mvncall_integrated_v5a.20130502.genotypes.vcf.gz
ALL.chr9.phase3_shapeit2_mvncall_integrated_v5a.20130502.genotypes.vcf.gz
ALL.chr10.phase3_shapeit2_mvncall_integrated_v5a.20130502.genotypes.vcf.gz
ALL.chr11.phase3_shapeit2_mvncall_integrated_v5a.20130502.genotypes.vcf.gz
ALL.chr12.phase3_shapeit2_mvncall_integrated_v5a.20130502.genotypes.vcf.gz
ALL.chr13.phase3_shapeit2_mvncall_integrated_v5a.20130502.genotypes.vcf.gz
ALL.chr14.phase3_shapeit2_mvncall_integrated_v5a.20130502.genotypes.vcf.gz
ALL.chr15.phase3_shapeit2_mvncall_integrated_v5a.20130502.genotypes.vcf.gz
ALL.chr16.phase3_shapeit2_mvncall_integrated_v5a.20130502.genotypes.vcf.gz
ALL.chr17.phase3_shapeit2_mvncall_integrated_v5a.20130502.genotypes.vcf.gz
ALL.chr18.phase3_shapeit2_mvncall_integrated_v5a.20130502.genotypes.vcf.gz
ALL.chr19.phase3_shapeit2_mvncall_integrated_v5a.20130502.genotypes.vcf.gz
ALL.chr20.phase3_shapeit2_mvncall_integrated_v5a.20130502.genotypes.vcf.gz
ALL.chr21.phase3_shapeit2_mvncall_integrated_v5a.20130502.genotypes.vcf.gz
ALL.chr22.phase3_shapeit2_mvncall_integrated_v5a.20130502.genotypes.vcf.gz
ALL.chrX.phase3_shapeit2_mvncall_integrated_v1b.20130502.genotypes.vcf.gz
ALL.chrY.phase3_integrated_v2a.20130502.genotypes.vcf.gz
ALL.chrMT.phase3_callmom-v0_4.20130502.genotypes.vcf.gz
```

#### 3.3 HRC

##### Description

The dataset describes a total of 40, 405, 505 variant sites at 27, 165 *H. sapiens* individuals.

It consists of 25 VCF files, corresponding to 22 autosomes and sex chromosomes (X_nonPAR, X_PAR1, X_PAR2).

The number of allowed FILTER/INFO/FORMAT fields (based on the headers of the files):
- 6 (1/4/1), including GT in FORMAT.

##### Source

The dataset is a reference panel from the Haplotype Reference Consortium [11].

The files were downloaded from the European Genom-phenom Archive (EGAS00000000029). The complete list of compressed VCF files used:

```
HRC.r1-1.EGA.GRCh37.chr1.haplotypes.noIBD.vcf.gz
HRC.r1-1.EGA.GRCh37.chr2.haplotypes.vcf.gz
HRC.r1-1.EGA.GRCh37.chr3.haplotypes.vcf.gz
HRC.r1-1.EGA.GRCh37.chr4.haplotypes.vcf.gz
HRC.r1-1.EGA.GRCh37.chr5.haplotypes.vcf.gz
HRC.r1-1.EGA.GRCh37.chr6.haplotypes.vcf.gz
HRC.r1-1.EGA.GRCh37.chr7.haplotypes.vcf.gz
HRC.r1-1.EGA.GRCh37.chr8.haplotypes.vcf.gz
HRC.r1-1.EGA.GRCh37.chr9.haplotypes.vcf.gz
HRC.r1-1.EGA.GRCh37.chr10.haplotypes.vcf.gz
HRC.r1-1.EGA.GRCh37.chr11.haplotypes.vcf.gz
HRC.r1-1.EGA.GRCh37.chr12.haplotypes.vcf.gz
HRC.r1-1.EGA.GRCh37.chr13.haplotypes.vcf.gz
HRC.r1-1.EGA.GRCh37.chr14.haplotypes.vcf.gz
HRC.r1-1.EGA.GRCh37.chr15.haplotypes.vcf.gz
HRC.r1-1.EGA.GRCh37.chr16.haplotypes.vcf.gz
HRC.r1-1.EGA.GRCh37.chr17.haplotypes.vcf.gz
```

HRC.r1-1.EGA.GRCh37.chr18.haplotypes.vcf.gz
HRC.r1-1.EGA.GRCh37.chr19.haplotypes.vcf.gz
HRC.r1-1.EGA.GRCh37.chr20.haplotypes.vcf.gz
HRC.r1-1.EGA.GRCh37.chr21.haplotypes.vcf.gz
HRC.r1-1.EGA.GRCh37.chr22.haplotypes.vcf.gz
HRC.r1-1.EGA.GRCh37.chrX_PAR1.haplotypes.vcf.gz
HRC.r1-1.EGA.GRCh37.chrX_PAR2.haplotypes.vcf.gz
HRC.r1-1.EGA.GRCh37.chrX_nonPAR.haplotypes.vcf.gz

#### 3.4 ExAC

##### Description

The dataset describes a total of 9,362,318 variant sites at exomes of *H. sapiens*.

It consists of 1 VCF files for 22 autosomes, X, Y and MT chromosomes, and contigs.

The number of allowed FILTER/INFO/FORMAT fields (based on the header of the file):

- 121 (17/97/7), but there are no FORMAT column (no FORMAT fields).

##### Source

The dataset is the aggregation of high-quality exome DNA sequence data for 60,706 individuals, generated as part of the Exome Aggregation Consortium (ExAC) [9].

The file were downloaded from:
https://storage.googleapis.com/gnomad-public/legacy/exac_browser/ExAC.r1.sites.vep.vcf.gz

#### 3.5 gnomAD2

##### Description

The dataset describes a total of 261,942,336 variant sites at *H. sapiens*.

It consists of 23 VCF files for 22 autosomes and X chromosome.

The number of allowed FILTER/INFO/FORMAT fields (based on the header of the file):

- all chromosomes: 572 (4/568/0).

##### Source

The dataset is a version 2.1.1 of the Genome Aggregation Database (gnomAD v2.1.1) [7]. It spans 15,708 whole-genome sequences from unrelated individuals.

The files were downloaded from:
https://gnomad.broadinstitute.org/downloads

The complete list of compressed VCF files used:

gnomad.genomes.r2.1.1.sites.1.vcf.bgz
gnomad.genomes.r2.1.1.sites.2.vcf.bgz
gnomad.genomes.r2.1.1.sites.3.vcf.bgz
gnomad.genomes.r2.1.1.sites.4.vcf.bgz
gnomad.genomes.r2.1.1.sites.5.vcf.bgz
gnomad.genomes.r2.1.1.sites.6.vcf.bgz
gnomad.genomes.r2.1.1.sites.7.vcf.bgz
gnomad.genomes.r2.1.1.sites.8.vcf.bgz
gnomad.genomes.r2.1.1.sites.9.vcf.bgz
gnomad.genomes.r2.1.1.sites.10.vcf.bgz
gnomad.genomes.r2.1.1.sites.11.vcf.bgz

gnomad.genomes.r2.1.1.sites.12.vcf.bgz
gnomad.genomes.r2.1.1.sites.13.vcf.bgz
gnomad.genomes.r2.1.1.sites.14.vcf.bgz
gnomad.genomes.r2.1.1.sites.15.vcf.bgz
gnomad.genomes.r2.1.1.sites.16.vcf.bgz
gnomad.genomes.r2.1.1.sites.17.vcf.bgz
gnomad.genomes.r2.1.1.sites.18.vcf.bgz
gnomad.genomes.r2.1.1.sites.19.vcf.bgz
gnomad.genomes.r2.1.1.sites.20.vcf.bgz
gnomad.genomes.r2.1.1.sites.21.vcf.bgz
gnomad.genomes.r2.1.1.sites.22.vcf.bgz
gnomad.genomes.r2.1.1.sites.X.vcf.bgz

#### 3.6 gnomAD3

##### Description

The dataset describes a total of $707,950,943$ variant sites at *H. sapiens*.

It consists of 24 VCF files for 22 autosomes, X and Y chromosomes.

The number of allowed FILTER/INFO/FORMAT fields (based on the header of the file):

- all chromosomes: 161 (4/157/0).

##### Source

The dataset is a version 3 of the Genome Aggregation Database (gnomAD v3) [7]. It spans 71,702 genomes from unrelated individuals.

The files were downloaded from:
`https://gnomad.broadinstitute.org/downloads`

The complete list of compressed VCF files used:

gnomad.genomes.r3.0.sites.chr1.vcf.bgz
gnomad.genomes.r3.0.sites.chr2.vcf.bgz
gnomad.genomes.r3.0.sites.chr3.vcf.bgz
gnomad.genomes.r3.0.sites.chr4.vcf.bgz
gnomad.genomes.r3.0.sites.chr5.vcf.bgz
gnomad.genomes.r3.0.sites.chr6.vcf.bgz
gnomad.genomes.r3.0.sites.chr7.vcf.bgz
gnomad.genomes.r3.0.sites.chr8.vcf.bgz
gnomad.genomes.r3.0.sites.chr9.vcf.bgz
gnomad.genomes.r3.0.sites.chr10.vcf.bgz
gnomad.genomes.r3.0.sites.chr11.vcf.bgz
gnomad.genomes.r3.0.sites.chr12.vcf.bgz
gnomad.genomes.r3.0.sites.chr13.vcf.bgz
gnomad.genomes.r3.0.sites.chr14.vcf.bgz
gnomad.genomes.r3.0.sites.chr15.vcf.bgz
gnomad.genomes.r3.0.sites.chr16.vcf.bgz
gnomad.genomes.r3.0.sites.chr17.vcf.bgz
gnomad.genomes.r3.0.sites.chr18.vcf.bgz
gnomad.genomes.r3.0.sites.chr19.vcf.bgz
gnomad.genomes.r3.0.sites.chr20.vcf.bgz
gnomad.genomes.r3.0.sites.chr21.vcf.bgz
gnomad.genomes.r3.0.sites.chr22.vcf.bgz

gnomad.genomes.r3.0.sites.chrX.vcf.bgz
gnomad.genomes.r3.0.sites.chrY.vcf.bgz

#### 3.7 SGDP

##### Description

The dataset describes 26 *H. sapiens* individuals with about 4 million variant sites for each (a total of 110, 291, 925 variant sites for the dataset).

It consists of 26 VCF files, each for a single individual.

The number of allowed FILTER/INFO/FORMAT fields (based on the header of the file):

- for each individual: 31 (2/23/6), including GT in FORMAT.

##### Source

The dataset is from the Simons Genome Diversity Project (SGDP) [6].

The files were downloaded and selected from 279 samples available here:
https://sharehost.hms.harvard.edu/genetics/reich_lab/sgdp/vcf_variants/vcfs.variants.public_samples.279samples.tar

The complete list of compressed VCF files used:

LP6005441-DNA_A01.annotated.nh2.variants.vcf.gz
LP6005441-DNA_A03.annotated.nh2.variants.vcf.gz
LP6005441-DNA_A04.annotated.nh2.variants.vcf.gz
LP6005441-DNA_A05.annotated.nh2.variants.vcf.gz
LP6005441-DNA_A06.annotated.nh2.variants.vcf.gz
LP6005441-DNA_A08.annotated.nh2.variants.vcf.gz
LP6005441-DNA_A10.annotated.nh2.variants.vcf.gz
LP6005441-DNA_A11.annotated.nh2.variants.vcf.gz
LP6005441-DNA_A12.annotated.nh2.variants.vcf.gz
LP6005441-DNA_B01.annotated.nh2.variants.vcf.gz
LP6005441-DNA_B02.annotated.nh2.variants.vcf.gz
LP6005441-DNA_B03.annotated.nh2.variants.vcf.gz
LP6005441-DNA_B04.annotated.nh2.variants.vcf.gz
LP6005441-DNA_B05.annotated.nh2.variants.vcf.gz
LP6005441-DNA_B06.annotated.nh2.variants.vcf.gz
LP6005441-DNA_B07.annotated.nh2.variants.vcf.gz
LP6005441-DNA_B08.annotated.nh2.variants.vcf.gz
LP6005441-DNA_B09.annotated.nh2.variants.vcf.gz
LP6005441-DNA_B10.annotated.nh2.variants.vcf.gz
LP6005441-DNA_B11.annotated.nh2.variants.vcf.gz
LP6005441-DNA_B12.annotated.nh2.variants.vcf.gz
LP6005441-DNA_C01.annotated.nh2.variants.vcf.gz
LP6005441-DNA_C02.annotated.nh2.variants.vcf.gz
LP6005441-DNA_C03.annotated.nh2.variants.vcf.gz
LP6005441-DNA_C05.annotated.nh2.variants.vcf.gz
LP6005441-DNA_C06.annotated.nh2.variants.vcf.gz

##### 3.8 Sheep

###### Description

The dataset describes a total of 42,438,841 variant sites at 99 *Ovis aries (sheep)* individuals.

It consists of 1 VCF file for 26 autosomes and sex chromosome X.

The number of allowed FILTER/INFO/FORMAT fields (based on the headers of the file):

- all chromosomes: 34 (1/26/7), including GT in FORMAT.

###### Source

The dataset is from the whole-genome sequencing of 99 sheep [13].

The file were downloaded from:
`ftp://parrot.genomics.cn/gigadb/pub/10.5524/100001_101000/100408/99sheep.fil.format2.vcf.gz`

### 3.9 AT

###### Description

The dataset describes a total of 119,146,348 variant sites at 1,135 *A. thaliana* samples.

It consists of 1 VCF file for 5 chromosomes.

The number of allowed FILTER/INFO/FORMAT fields (based on the headers of the file):

- all chromosomes: 7 (3/1/3), including GT in FORMAT..

###### Source

The dataset is from the 1001 Genomes Project [1].

The file were downloaded from:
`https://1001genomes.org/data/GMI-MPI/releases/v3.1/1001genomes_snp-short-indel_with_tair10_only_ACGTN.vcf.gz`

###### Comments

The file was split into 5 smaller parts, each for a single chromosome. The header of the files were modified, to comply with the VCF specification, i.e.:

- The missing contigs (chromosomes) description were added:
```
##contig=<ID=1>
##contig=<ID=2>
##contig=<ID=3>
##contig=<ID=4>
##contig=<ID=5>
```

##### 3.10 Rice

###### Description

The dataset describes 6 *Oryza sativa L.(rice)* samples with about 409 million variant sites for each (a total of 20,617,610,298 variant sites for the dataset).

It consists of 6 gVCF files, each for a single sample.

The number of allowed FILTER/INFO/FORMAT fields (based on the header of the file):

- for each individual: 27 (2/20/5), including GT in FORMAT.

##### Source

The dataset is from the The 3000 Rice Genome Project [15].

The files were downloaded and selected from 2466 samples available here:
`http://iric.irri.org/resources/3000-genomes-project`

The complete list of compressed VCF files used:

`IRIS_313-10000.snp.vcf.gz`
`IRIS_313-10001.snp.vcf.gz`
`IRIS_313-10002.snp.vcf.gz`
`IRIS_313-10007.snp.vcf.gz`
`IRIS_313-10010.snp.vcf.gz`
`IRIS_313-10014.snp.vcf.gz`

#### 3.11 Tomato

##### Description

The dataset describes a total of 11,620,517 variant sites at 361 *S. lycopersicum (tomato)* individuals.

It consists of 1 VCF file for 12 chromosomes.

The number of allowed FILTER/INFO/FORMAT fields (based on the headers of the file):
- all chromosomes: 13 (1/9/3), including GT in FORMAT.

##### Source

The dataset is from the AGIS Tomato 360 Resequencing Project [10].

The file were downloaded from:
`ftp://ftp.solgenomics.net/genomes/tomato_360//360_merged_2.50.vcf.gz`

##### Comments

The header of the `360_merged_2.50.vcf` file was modified, to comply with the VCF specification, i.e.:

- The missing definition of "AC1" field was added, assuming type String (INFO field): `##INFO=<ID=AC1,Number=1,Type=String>`
- The missing definition of "GT" field was added, assuming type Flag (INFO field): `##INFO=<ID=GT,Number=1,Type=Flag>`
- The missing definition of "0|1" field was added, assuming type String (FORMAT field): `##FORMAT=<ID=0|1,Number=1,Type=String>`
- The missing definition of "1|1" field was added, assuming type String (FORMAT field): `##FORMAT=<ID=1|1,Number=1,Type=String>`
- The missing contigs (chromosomes) description were added:
  `##contig=<ID=SL2.50ch00>`
  `##contig=<ID=SL2.50ch01>`
  `##contig=<ID=SL2.50ch02>`
  `##contig=<ID=SL2.50ch03>`
  `##contig=<ID=SL2.50ch04>`
  `##contig=<ID=SL2.50ch05>`
  `##contig=<ID=SL2.50ch06>`
  `##contig=<ID=SL2.50ch07>`
  `##contig=<ID=SL2.50ch08>`
  `##contig=<ID=SL2.50ch09>`

```
##contig=<ID=SL2.50ch10>
##contig=<ID=SL2.50ch11>
##contig=<ID=SL2.50ch12>
```

#### 4 Environment

The computer used in tests was of the following configuration:

- 2 Intel Xeon E5-2670 v3 CPUs, 12 double-threaded cores per CPU, each clocked at 2.3 GHz,
- 128 GiB RAM.
- 2 HDDs of size 6 TiB each in RAID-0 configuration, `hdparm -t` reported buffered read speed 411 MB/s.

For compilation we used G++ v. 9.3.1. The machine was running CentOS 7.

### 5 Additional results

Additional results are given in the Supplementary Worksheet.
